# High viral abundance and low diversity are associated with increased CRISPR-Cas prevalence across microbial ecosystems

**DOI:** 10.1101/2021.06.24.449667

**Authors:** S. Meaden, A. Biswas, K. Arkhipova, S. E. Morales, B. E. Dutilh, E. R. Westra, P. C. Fineran

**Author notes:** These authors contributed equally to this work (S. Meaden, A. Biswas). These authors share senior authorship (S. E. Morales, B. E. Dutilh, E. R. Westra, P. C. Fineran). Correspondence: Sean Meaden.

## Abstract

CRISPR-Cas are adaptive immune systems that protect their hosts against viruses and other parasitic mobile genetic elements. Consequently, selection from viruses and other genetic parasites is often assumed to drive the acquisition and maintenance of these immune systems in nature, but this remains untested. Here, we analyse the abundance of CRISPR arrays in natural environments using metagenomic datasets from 332 terrestrial, aquatic and host-associated ecosystems. For each metagenome we quantified viral abundance and levels of viral community diversity to test whether these variables can explain variation in CRISPR-Cas abundance across ecosystems. We find a strong positive correlation between CRISPR-Cas abundance and viral abundance. In addition, when controlling for differences in viral abundance, we found that the CRISPR-Cas systems are more abundant when viral diversity is low. We also found differences in relative CRISPR-Cas abundance among environments, with environmental classification explaining ∼24% of variation in CRISPR-Cas abundance. However, the correlations with viral abundance and diversity are broadly consistent across diverse natural environments. These results indicate that viral abundance and diversity are major ecological factors that drive the selection and maintenance of CRISPR-Cas in microbial ecosystems.

**Significance statement:** Numerous studies demonstrate that CRISPR-Cas immune systems can provide defence against bacteriophage and archaeal viruses, yet little is known about the ecological conditions where CRISPR-Cas immunity is favoured. Moreover, our knowledge is largely confined to laboratory studies and it is unknown if viruses are a key selective driver of CRISPR-Cas in nature. Using metagenomic data from diverse environments we find that both viral abundance and the abundance of CRISPR-Cas immune systems correlate positively across most environments. Furthermore, CRISPR-Cas systems are more prevalent when viral diversity is low. These results extend previous theoretical work by demonstrating that viruses are a key driver of selection of CRISPR-Cas immune systems across many natural ecosystems.

## Introduction

CRISPR-Cas is a sophisticated immune system of bacteria and archaea that is widespread across diverse prokaryotic taxa [1]. Anecdotal evidence suggests that CRISPR-Cas systems are unevenly distributed across environments and species. For example, culturable bacteria seem to have higher frequencies of CRISPR-Cas than non-culturable lineages [2], and extremophiles tend to have higher CRISPR-Cas content and longer CRISPR arrays than mesophiles [3, 4]. Given that CRISPR-Cas, like most defence system genes, are frequently lost and gained, this strongly suggests that CRISPR-Cas is advantageous under particular environmental conditions [5]. However, environmental parameters that determine the benefits of CRISPR-Cas remain unclear. *In vitro* studies suggest that the density of phages is a key fitness determinant of CRISPR-Cas immune systems [6], as high phage density favours the evolution of surface-based resistance over CRISPR-Cas. However, whether phage density explains the distribution of CRISPR-Cas in natural environments remains unexplored. Moreover, theory and laboratory experiments suggest that environments with genetically diverse phage populations may favour organisms with generalist defences over those containing the specific CRISPR-Cas system [7].

While these questions have been explored using theory and controlled laboratory-based experiments, they are rarely assessed using environmental samples [8–11]. The increasing availability of metagenomic data from a range of environments now provides an opportunity to test these hypotheses. Here, we quantify CRISPR array abundance and diversity across metagenomic samples that span a range of terrestrial, aquatic and host-associated ecosystems. We combine these data with measures of viral abundance and diversity to test to what extent these variables explain the distribution of CRISPR-Cas systems in natural environments. Across different types of environments, we consistently find a positive correlation between CRISPR-Cas abundance and viral abundance and a negative correlation with viral diversity. Taken together, these results imply that CRISPR-Cas is most beneficial in environments with high viral abundance but low diversity and that these are important drivers of the prevalence of CRISPR-Cas immune systems in the environment.

## Results

### Variation in CRISPR-Cas abundance is partially explained by viral abundance

While it is well-established that CRISPR-Cas immune systems can protect bacteria and archaea against viral infections under *in vitro* laboratory conditions, it remains unclear how important these genetic parasites are as a selective force for the maintenance of CRISPR-Cas systems in nature [12]. To assess the role of viruses as a selective force for CRISPR-Cas, we first compiled a dataset of 332 metagenomes and quantified the abundance of CRISPR-Cas systems and viruses in each sample. We note that our analyses uses all contigs classified as viral, and while the vast majority of these are of bacteriophage origin, we refer to these simply as viral for consistency (benchmarking analyses of our classifier vs. existing tools is available in the SI). These metagenomes vary in both CRISPR-Cas and viral abundance and therefore, provided a suitable dataset to test the hypothesis that viral abundance drives selection for CRISPR-Cas (Fig. 1). We found a positive correlation between viral abundance and the abundance of CRISPR-Cas systems (GLM, F_1,295_ = 77.78, p < 0.0001, Fig. 1), with viral abundance explaining around 20% of the observed variation in CRISPR-Cas abundance (R^2^ = 0.209). We obtained qualitatively the same result when we included archaeal abundance in our model, which typically carry more CRISPR-Cas immune systems than bacteria [1]. These results strongly suggest that viruses are a fundamental selective force for the maintenance of CRISPR-Cas across diverse environmental conditions.

**Figure 1.**
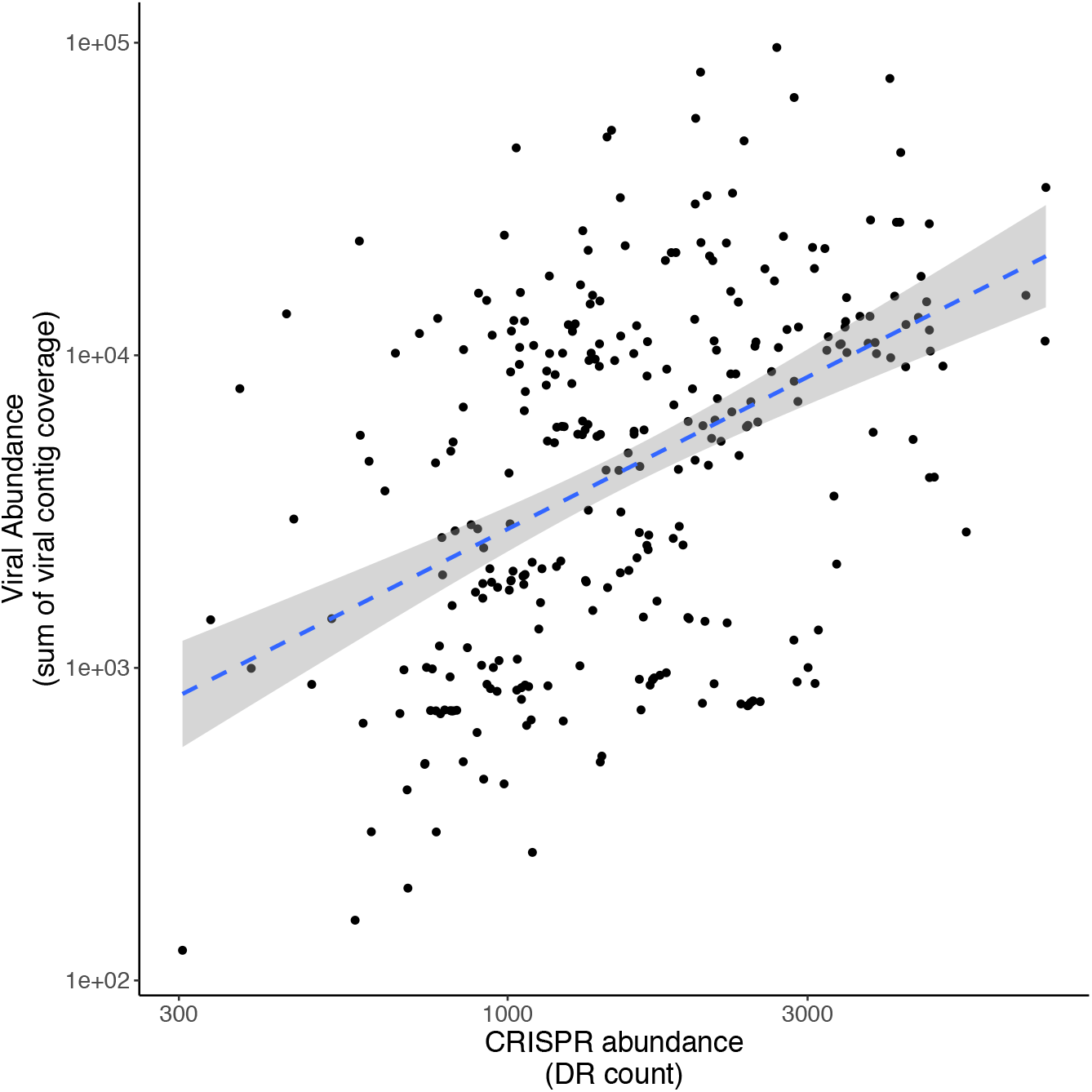
CRISPR abundance positively correlates with viral abundance. Correlation between relative viral abundance and the read count (per million) of metagenomic reads that mapped to CRISPR array repeats across all samples. The dashed line represents the linear model fit and shaded area represents 95% confidence interval (p-value < 0.0001 and R^2^ = 0.21).

### Environmental conditions influence CRISPR-Cas abundance

In addition to viruses being a selective force for CRISPR-Cas, ecological factors may determine when CRISPR-Cas is beneficial and therefore, CRISPR-Cas abundance may vary across different natural environments [13]. We therefore grouped samples into ecologically meaningful categories, using the Earth Microbiome Project’s sample ontology (EMPO). This framework is structured to capture two major environmental axes on which bacterial community composition orient, namely host-association and salinity [14]. Level 1 of the ontology classifies samples as host-associated or free living, level 2 classifies samples as saline or non-saline, or animal or plant-associated and level 3 describes microbial environments that can be grouped into levels 1 and 2 hierarchically (Fig. 2A [14]). These EMPO classifications highlighted the varied CRISPR-Cas and viral abundance in these different environments (Fig. 2 and Fig. S2). Using this EMPO classification, we observed substantial variation in CRISPR-Cas abundance, both within and between environment types (Fig. 2). For example, host-associated communities had a greater prevalence of CRISPR-Cas than free-living environments (GLM, F_1,330_ = 7.01, p = 0.008), although this classification only explained around 2% of the variation in CRISPR-Cas abundance. In contrast, more fine scaled classification of environments (such as gut, saline sediment etc. as per EMPO level 3 classification, Fig. 2C), explained 24% of the variation in CRISPR-Cas abundance. Taken together, these results suggest that in addition to viruses as a key selective force for the maintenance of CRISPR-Cas, there are substantial differences in CRISPR-Cas abundance among natural environments.

**Figure 2.**
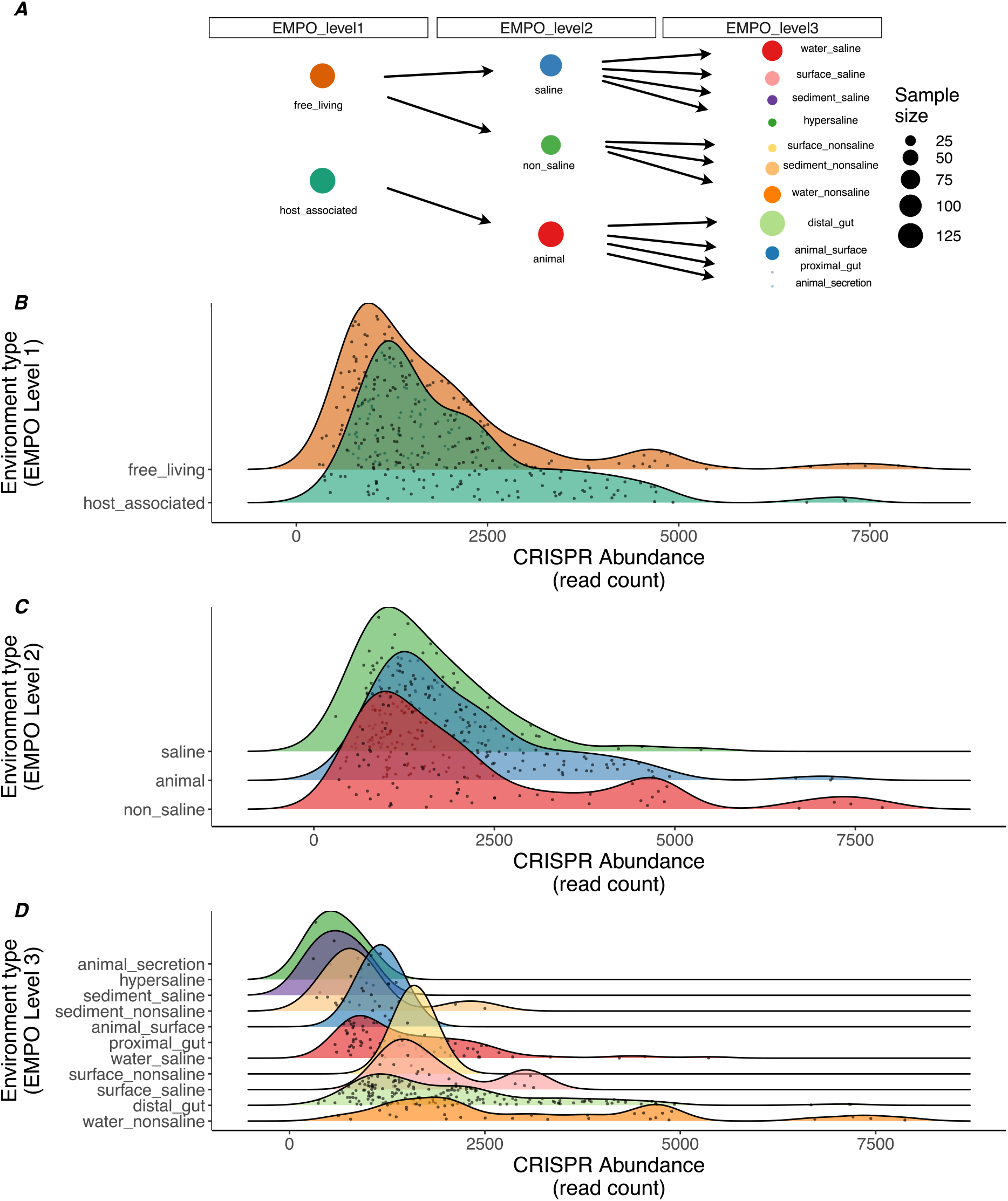
CRISPR abundance varies by environment. Distributions of metagenomic read counts that mapped to CRISPR arrays (read count per million that mapped to a CRISPR array predicted by CRISPRDetect v.3 from assembled contigs) grouped by environmental classification. (A) Sample sizes and ontology is shown in panel A. Samples are grouped using the Earth Microbiome Project Ontology (EMPO) at level 1 (B), 2 (C) or 3 (D).

### Microbial community composition explains some variation in CRISPR-Cas abundance

Although we found effects of viral abundance and environmental classification on CRISPR-Cas abundance, it is plausible that these effects may be driven by differences in the microbial community composition as CRISPR-Cas prevalence can differ among taxa [1, 2]. We examined whether the variation in CRISPR-Cas abundance across these metagenomes might be affected microbial community composition. We found a weak but significant relationship between CRISPR-Cas abundance and class-level community composition (PERMANOVA, F_1, 313_ = 12.4, p < 0.001, R^2^ = 0.04, permutations = 9999, distance metric = Bray-Curtis). We next used clustering analyses to assess how well the EMPO framework levels grouped our samples based on community composition and extracted the taxa that best described differences among samples (Fig. S3A). In addition, we fitted CRISPR-Cas abundance to this ordination to identify ‘hotspots’ of samples enriched with CRISPR-Cas (Fig. S3B). With this approach, we identified multiple groups of samples with high CRISPR-Cas abundance, supporting the notion that microbial phylogeny alone only explains a limited amount of variation in CRISPR-Cas abundance. Additional factors may contribute to the differences in CRISPR-Cas abundance across environment types. Furthermore, high rates of horizontal gene transfer (HGT) across taxa, coupled with frequent gain or loss of CRISPR-Cas, will likely reduce phylogenetic signal.

As an additional test of the influence of phylogenetic effects on our results, we assessed the impact of including archaeal abundance in our analyses. Archaea have previously been shown to be enriched for CRISPR-Cas systems [1]; therefore, inclusion of archaeal abundance in our model should control for this effect. When we repeated our analysis of the correlation between CRISPR-Cas abundance and viral abundance, this time including the abundance of archaea in each sample as a covariate, we found no qualitative difference in result. Taken with the community composition analysis, these results suggests that the influence of phylogeny on our results is relatively small compared to the effect of viral abundance on the prevalence of CRISPR-Cas immune systems within a microbiome.

### CRISPR-Cas and viral abundance correlate across diverse environments

To explore to what extent the observed variation in CRISPR-Cas abundance within and between environment types is driven by variation in viral abundance, we grouped the samples by EMPO classification and quantified viral abundance for each environment type. Similarly, to the distributions of CRISPR-Cas abundance, we found substantial variation in viral abundance across environment types (Fig. S2). The higher-level classifications, EMPO level 1 and 2, explained 30% and 31% of the variation observed respectively, and the finer scale EMPO level 3 explained 34% of this variation in viral abundance (GLM, F_1,285_ = 13.14, p < 0.0001, R^2^ = 0.34, Fig. S2C). The relatively minor difference between classification levels suggests that the primary predictive power comes from EMPO level 1, with host-associated samples having a greater density of viruses than free-living samples. Overall, we found that the type of environment significantly predicts the abundance of viruses present, but that fine scale classification adds little predictive power relative to high-level classification.

Although we found a significant correlation between viral abundance and CRISPR-Cas abundance, it remained unclear whether this relationship was consistent across environments. We therefore assessed whether the strength of this relationship was constant among each of our EMPO classification levels, which we modelled as an interaction in a multiple regression analysis. Strikingly, we found significant interaction effects at all EMPO levels, suggesting that the nature of the relationship between viral abundance and CRISPR-Cas abundance is, at least partly, dependent on additional environmental conditions (Table 1, Fig. 3). This was further validated by post-hoc testing of the correlation between CRISPR-Cas abundance and viral abundance at each individual environment type (Table S3). In this case we found a consistent positive relationship at EMPO levels 1 and 2, but more varied results at level 3 (Fig. 3), suggesting additional ecological factors may be playing a role in some environments. For example, when taking all EMPO level 3 classifications with more than 10 observations per group, non-saline sediments show a strong positive correlation between CRISPR-Cas abundance and viral abundance (adjusted p-value < 0.001, Pearson correlation = 0.71, n = 15). In contrast, non-saline water environmental samples show no significant correlation between CRISPR-Cas abundance and viral abundance (adjusted p-value = 1, Pearson correlation = -0.18, n = 34). All correlations can be found in Table S4. Taken together, these results indicate that viral abundance typically correlates positively with CRISPR-Cas abundance, but that the strength of this relationship is dependent on the particular environment.

**Table 1.** Interaction effects between virus abundance and CRISPR across environments. Linear models used to test the relationship between CRISPR abundance (DR count) and viral abundance at each EMPO classification level.

| Resid. Df | Resid. Dev | df | Deviance | F-value | P-value | AIC | R<sup>2</sup> | Model formula |
| --- | --- | --- | --- | --- | --- | --- | --- | --- |
| **294** | 15.345 | -1 | -0.288 | 5.595 | 0.019 | -32.758 | 0.224 | glm(log10(DR_count) ~ log10(viral_abund) * EMPO_level1 |
| **293** | 15.308 | -2 | -0.789 | 7.908 | <0.001 | -39.572 | 0.252 | glm(log10(DR_count) ~ log10(viral_abund) * EMPO_level2 |
| **284** | 12.074 | -9 | -1.086 | 3.02 | 0.002 | -90.349 | 0.434 | glm(log10(DR_count) ~ log10(viral_abund) * EMPO_level3 |

**Figure 3.**
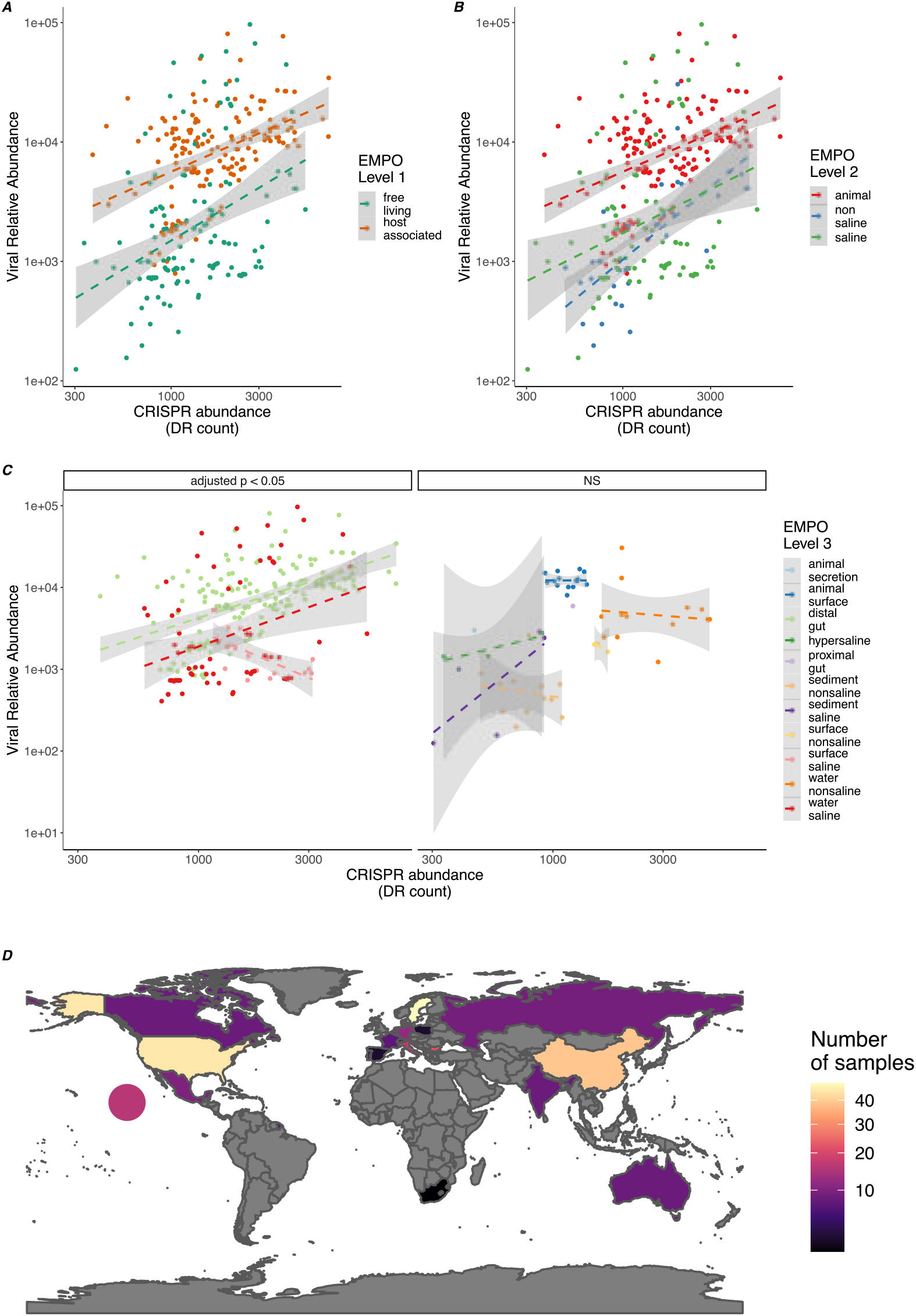
CRISPR abundance positively correlates with viral abundance across environments. Correlations between relative viral abundance and the read count (per million) of metagenomic reads that mapped to CRISPR array repeats per environment type. Environments are categorised according to the Earth Microbiome Project Ontology at level 1 (A) or 2 (B). Samples grouped at EMPO level 3 (C) are divided into significant correlations or non-significant correlations (NS). Dashed lines represent linear model fits and shaded areas represent 95% confidence intervals. (D) The number of samples collected in each country with the circle representing samples collected in the Pacific Ocean.

### Virus diversity negatively correlates with CRISPR-Cas abundance

Both theory and *in vitro* experiments predict that low viral genetic diversity is also an important determinant of the benefits of a CRISPR-Cas immunity [7–9, 11]. This theory suggests that excess viral sequence diversity prevents the acquisition of sufficient spacer diversity to protect against the many different viruses. To test this prediction, we quantified viral diversity for each environment type and examined if this correlated with CRISPR-Cas abundance. However, this analysis may be confounded by correlations between viral diversity and viral abundance. Indeed, in our dataset viral diversity was strongly correlated with viral abundance (GLM, F_1,295_ = 208.6, p < 0.0001, R^2^ = 0.41, Fig. S4). We therefore normalized the viral diversity scores by viral abundance for each sample. We then tested the correlation between CRISPR-Cas abundance and normalized viral diversity. In agreement with theory, we found a negative correlation between CRISPR-Cas abundance and normalized viral diversity for all viral diversity metrics used (Fig. 4, Table 2). The metrics used spanned multiple levels, with richness, evenness and Shannon’s index describing inter-population diversity. By contrast, Nei’s diversity metric describes the intra-population genetic variation. Together these results suggest that CRISPR-Cas is most effective when viral diversity is low, which supports that CRISPR-Cas immunity relies on sequence identity between spacer sequences and the viral protospacer sequence and array sizes are finite.

**Table 2.**
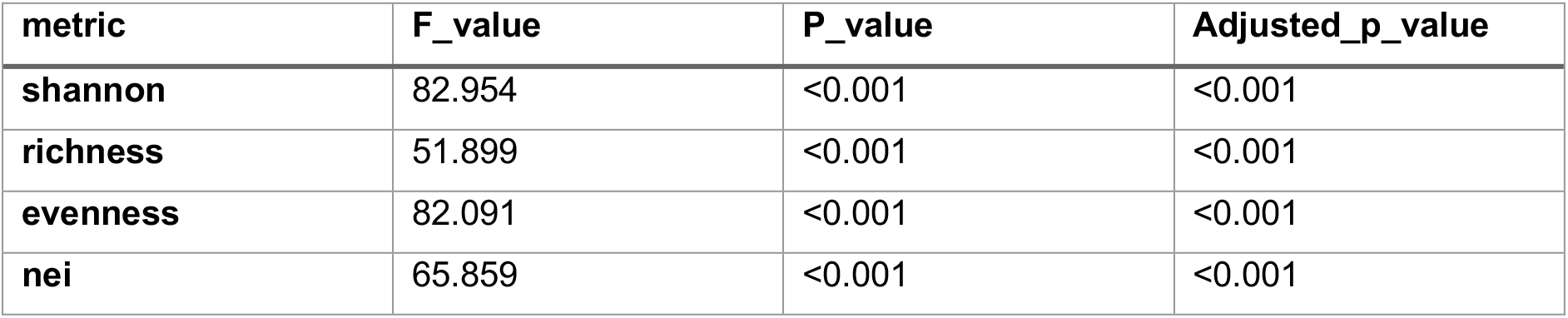
Associations between normalized viral diversity metrics and CRISPR abundance. Results from F-tests using linear models with Bonferroni adjusted p-values for the number of metrics tested (n = 4).

| metric | F_value | P_value | Adjusted_p_value |
| --- | --- | --- | --- |
| shannon | 82.954 | <0.001 | <0.001 |
| richness | 51.899 | <0.001 | <0.001 |
| evenness | 82.091 | <0.001 | <0.001 |
| nei | 65.859 | <0.001 | <0.001 |

**Figure 4.**
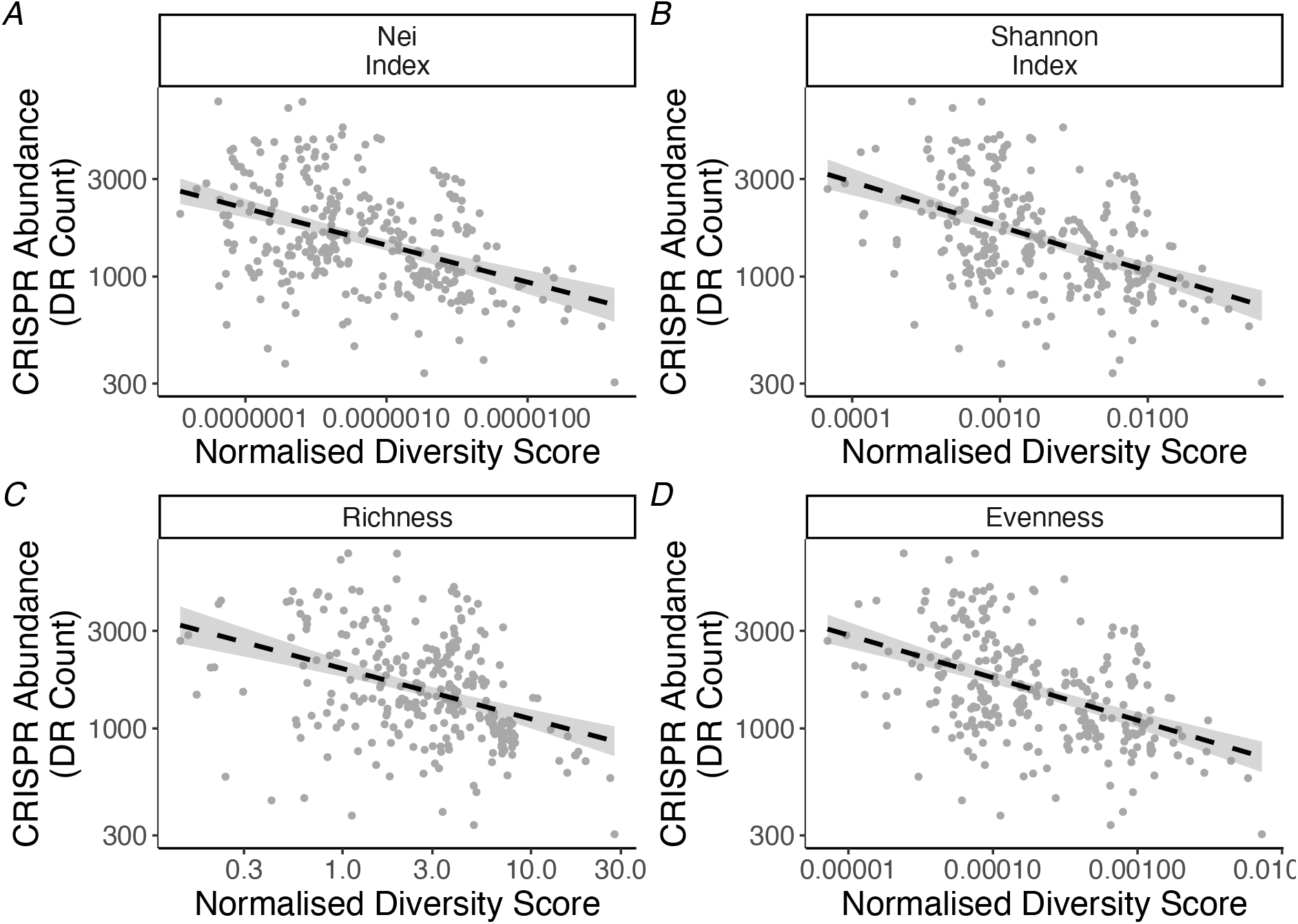
CRISPR abundance negatively correlates with normalised viral diversity metrics. Correlations between viral diversity (normalized by viral load per sample) and CRISPR abundance (reads per million that map to CRISPR arrays). Panels represent Nei’s diversity index (A), Shannon’s index (B), contig richness (C), contig evenness (D). Panel A represents intra-contig viral diversity while panels B, C and D represent inter-contig viral diversity.

## Discussion

Despite recent studies suggesting that CRISPR-Cas abundance varies across natural environments, such as soil [15] and the human microbiome [16], the ecological factors that drive variation in CRISPR-Cas prevalence across natural microbial communities remained unclear [12]. Furthermore, the extent of this variation across a much wider range of environments remained unexplored. We addressed this gap by using metagenomic data to quantify CRISPR array abundance within each metagenome and linked these data to the associated viral community present. We identified two key factors that predict CRISPR-Cas abundance: viral abundance and viral diversity. These factors are likely of primary importance as the observed correlations are consistent across a broad range of environments. Taken together, our results show that high viral abundance and low diversity are major drivers of the selection and maintenance CRISPR-Cas systems in nature.

There are likely factors in addition to viral abundance and diversity that contribute to CRISPR-Cas abundance in the environment because the correlations had relatively low R^2^ values (20% for abundance and 22% for normalized diversity). For example, many alternative phage defence systems have been described and, much like CRISPR-Cas, show scattered distributions even in closely related bacterial strains [17]. The interplay and redundancy between different phage defence systems is poorly characterised and may also contribute to CRISPR-Cas distributions in different environments. Understanding the environmental parameters that different defences will be crucial future research. For example, we see greater CRISPR-Cas abundance in host-associated samples over free-living samples, but it is unknown if alternative defence mechanisms are favoured in these free-living samples, or if there are fewer defences overall.

Regarding CRISPR-Cas, multiple environmental parameters have been predicted to interact negatively with these systems. For example, aerobicity showed a negative association with CRISPR-Cas prevalence during a modelling analysis of bacterial traits [13], possibly due to an incompatibility between the requirement for non-homologous end join repair (NHEJ) in aerobic respiration and type II CRISPR-Cas sytems [18]. More generally, intracellular defences may be favoured over surface receptor modifications under certain environmental conditions, as these have been shown to be subject to trade-offs with both biotic and abiotic factors [19–21]. In addition, recent work has demonstrated that regulation of phage defences, including CRISPR-Cas, is mediated by environmental conditions [22]. We also cannot exclude the role of plasmids in our analyses, as a recent longitudinal study found plasmids were targeted by CRISPR-Cas systems at 5-times the rate of phages [23]. Overall, our results demonstrate that phage-mediated selection is a major driver of CRISPR-Cas prevalence, but additional biotic and abiotic complexity likely shape the strength of this relationship.

Previous theoretical models predicted that CRISPR-Cas will be less favourable in dense and diverse viral communities [8, 9]. Above a threshold of phage genetic diversity CRISPR-Cas becomes ineffective and is lost due an associated fitness cost and this threshold is reached more often in large viral populations [8]. While these predictions are intuitive, our results suggest that while low viral diversity does indeed favour CRISPR-Cas, low viral density does not. It is possible that viral abundance rarely reaches levels in nature that are sufficient to preclude an effective CRISPR-Cas response, even if such densities are readily achievable in laboratory experiments [10, 12]. Future work may reveal the ecological differences between CRISPR-Cas types, as different types are likely to coevolve with viruses in fundamentally different ways [24].

Genomic evidence demonstrate that CRISPR-Cas systems are frequently acquired and lost [25, 26] and empirical studies that show that they can be mobilized through HGT [27, 28]. Notably, in a previous longitudinal study, CRISPR-Cas abundance increased through time even in phyla decreasing in abundance under soil warming conditions [15], again suggesting high mobility and positive selection for CRISPR-Cas immunity. Consistent with high rates of HGT of CRISPR-Cas, our results demonstrate that while phylogeny can influence the CRISPR-Cas repertoire, it is not the primary driver of selection in nature.

In summary, by quantifying the role of the viral community in shaping CRISPR-Cas abundance in complex, diverse natural communities we found that high viral abundance, but low diversity, drives the selection and maintenance of CRISPR-Cas across a range of environments. Future work that embraces both the abiotic and biotic complexity of natural systems is required to further understand the prevalence of CRISPR-Cas.

## Methods

### Computational pipeline

An overview of the computational pipeline is provided in Figure S1.

### Collection of samples and assembly

Paired-end metagenomic libraries (sequenced with Illumina platform) were downloaded from the NCBI SRA database (Table S1). Libraries were processed with BBMap suite of tools for error correction [12] and for each metagenomic library a representative FASTA file was created by combining both the merged and unmerged reads. A total of 332 libraries (each library containing at least 1 million reads with a minimum length of 100 nucleotides (nt) and insert size >=150) were selected to represent a wide range of biome diversity (table S1). Libraries were assembled using MegaHit (version 1.1.3) [13] with default parameters and contigs with minimum length 200nt were retained. Contigs were classified as archaea, bacteria or virus using a purpose-built classification tool: MIUMS (Microbial Identification Using Marker Sequence, https://github.com/ambarishbiswas/miums_v1.0). MIUMS is designed to classify contigs based on a reference database containing protein sequence fragments highly specific to bacteria, archaea, viruses. Full details of MIUMS reference database construction and prediction process are given in the supplementary information. Each metagenome was then subsampled to 1 million randomly selected reads with a minimum length of 100 nt.

### Generation of archaeal and bacterial abundance tables

Subsampled reads were screened using metaxa2 [14] (GSU parameters: -g ssu -f f, LSU parameters: -g lsu -f f) to generate a table of bacterial and archaeal abundances. Both the LSU and SSU based methods were used. A reference sequence database was made from contigs classified as viral by MIUMS. Subsampled reads were then mapped to the assembled contigs using Magic-BLAST [15] (parameters: -no_unaligned -no_query_id_trim -perc_identity 95 -outfmt tabular). Reads with a minimum of 95% sequence identity and coverage were used.

### CRISPR array identification

Accurately identifying CRISPR arrays in metagenomic data is challenging for a number of reasons. Firstly, a large proportion of the direct repeats (DRs) identified from metagenomic contigs often show little sequence similarity to the CRISPR repeats found in published genomes and lack an isolated representative [2]. Secondly, CRISPR arrays found in metagenomic reads are generally short (i.e. < 3 DRs) and often missing one or both flanking regions. To overcome these issues we combined information on existing, published genomes and their CRISPR arrays as well *de novo* extraction of putative CRISPR arrays from our assembled contigs (Fig. S1). A database of metagenomic CRISPR arrays was first constructed by processing all assembled contigs with CRISPRDetect version 3 (CRISPRDetect3, https://github.com/ambarishbiswas/CRISPRDetect_3.0). CRISPRDetect version 3 was also modified to allow prediction of shorter CRISPRs (e.g. partial/broken CRISPRs with as little as 1.5 repeats). A higher CRISPR likelihood score cut-off of 4.5 was used instead of the default score cut-off of 3 to reduce potential non-CRISPR arrays. CRISPRDetect [16] uses several CRISPR elements (e.g. repeats, spacers, cas genes, AT composition of flanking regions etc.) from published genomes to identify and separate true CRISPRs from other genomic repeats. In this study, a modified version of the CRISPRDetect tool was used, which uses a reference repeat database created using the cluster representative DRs from the metagenomic contigs as well as DRs found in published genomes. Predicted CRISPR arrays were checked to ensure that the total array degeneracy (i.e. number of insertion, deletion, mutation or presence of Ns in the array) was less than the total number of DRs in the array, which resulted in 51395 CRISPR arrays. Direct repeat sequences (23 to 60 nt) were extracted and clustered with cd-hit-est (parameters: -n 3 -c 0.90 -aL 0.90 -aS 0.90) [17], which resulted in 33745 repeat clusters. A vast majority of these clusters (i.e. 22808) consisted of a single DR. This low level of redundancy was important to ensure the successful subsequent mapping of reads to DRs.

Subsampled reads were then screened against this database using metaCRISPRDetect (https://github.com/ambarishbiswas/metaCRISPRDetect_1.0), which supports rapid identification of CRISPR arrays in short reads using user-provided reference repeat database as an extension of CRISPRDetect [16]. Arrays with a likelihood score > 3 were added to the existing CRISPR reference database. Subsampled reads were then mapped to the reference database using blastn [18] [parameters: -task blastn-short with default parameters]. Abundance tables were generated for all spacers and direct repeat sequences by tallying the number of reads that mapped to a given DR with 100% sequence identity and coverage.

### Identifying potential false positive CRISPRs

CRISPRs predicted from metagenomes are generally short, often incomplete and missing flanking regions which makes it hard to distinguish true CRISPRs from other genomic repeats. To measure how many of our identified arrays occurred in known prokaryotic genomes we compared the metagenomic CRISPR repeats against CRISPRs found in RefSeq and GenBank prokaryotic sequences (sequences published before September 9, 2019) using NCBI blast (parameters: -task blastn -word_size 11 -dust no -culling_limit 1 -num_alignments 1) [19]. Metagenomic DRs with >=90% identity and >=90% sequence coverage against RefSeq or GenBank DRs were considered as a positive match. Out of the 7562 repeat clusters 1649 were found in CRISPRs predicted from RefSeq or GenBank prokaryotic sequences. Similarly, the metagenomic repeats were screened against an *in silico* generated set of eukaryotic reads from 6 eukaryotic reference genomes (Table S2). Ten thousand 250bp paired reads were generated from each reference genome using WGSIM (https://github.com/lh3/wgsim). Eukaryotic reads were subsampled to an equal depth to remove differences due to genome size. Out of the 7562 DR clusters a total of 168 repeat clusters were found in the eukaryotic reads suggesting a high level of eukaryotic sequence contamination may increase the false positive rate of our analysis. We therefore removed samples where > 10% of reads were classified as of eukaryotic origin.

### Virome analysis

To assess the diversity of viruses in the different environments, we analysed all MIUMS classified contigs from the 332 metagenomes. Across all datasets, we identified 2583 archaeal prophages, 375179 bacterial prophages, 1218279 bacteriophages, and 174792 archaeal viruses. Per metagenome, the sub-sampled reads used for the CRISPR quantification analysis were mapped to the total set of all viral contigs with bwa mem [20]. To quantify the genetic diversity of the viral community, we calculated the richness as the total number of detected viral contigs, plus the evenness and Shannon’s diversity index based on relative abundances calculated from the mean depth of coverage of all detected viral contigs. The intra-population diversity (micro-diversity) was calculated as the mean heterozygosity of viruses in the community by averaging Nei’s per-nucleotide diversity index across all detected contigs [21].

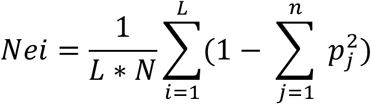

where p_j_ is the frequency of allele *j* at the position *i* of a contig of the length L, N is the number of viruses in a dataset. Single nucleotide variability was assessed with VarScan software [22].

Viral abundance was calculated by first collecting the average depth values for all viral contigs in each sample using the ‘jgi_summarize_bam_contig_depths’ script from the Metabat package [23]. These depth scores were then summed per sample to give an approximation of overall viral abundance relative to bacterial abundance.

### Statistical analysis

For each sample, the number of reads that mapped to a direct repeat were counted to give a measure of CRISPR array abundance per sample. Earth Microbiome Project Ontology levels were assigned using the framework previously described [24](Fig 1D) based on the associated metadata from the NCBI short read archive (SRA). In the case of bioreactor samples, a literature search was conducted to identify the original material described in the associated study (see Table S1).

General linear models (GLM) were constructed for each of the reported correlations. In each case log10 transformations were applied to conform to model assumptions. Checks of model residuals were performed to assess model fit. Significance was determined using F-tests between null models and those containing the variable or interaction of interest.

### Microbial community assessment

CRISPR is more common in archaea. Therefore, in order to minimise any phylogenetic effects deriving from high archaeal abundances in samples we estimated the number of archaeal reads in the subsampled reads using metaxa2 [14] and converted this to relative abundance per sample. We then included this value as a fixed effect in additional GLMs to check the influence of archaeal abundance on our results. For additional phylogenetic assessments, reads classified as non-prokaryotic were removed and relative abundances were generated using the output from metaxa2 in order to assess effects of community composition. Permutational ANOVAs were performed on species abundance matrices using Bray-Curtis dissimilarity and CRISPR abundance as explanatory variable with 9999 permutations, using the function ‘adonis’ from ‘vegan’ package [25]. Visualisation of clustering analyses, at the class level, was performed using non-metric multidimensional scaling (NMDS) with Bray-Curtis dissimilarity through the ‘metaMDS’ function in ‘vegan’ [25]. Smooth surfaces were fit to these points via a generalized additive model (GAM), using either CRISPR abundance as the explanatory term, with the ‘ordisurf’ function in ‘vegan’[25].

## Acknowledgments

This work was funded by a grant from the European Research Council under the European Union’s Horizon 2020 research and innovation programme (ERC-STG-2016-714478 to E.R.W.) E.R.W. was further supported by a NERC Independent Research Fellowship (NE/M018350/1). S.M. received funding from the European Union’s Horizon 2020 research and innovation programme under the Marie Skłodowska-Curie grant agreement No. 842656. B.E.D was supported by the Netherlands Organization for Scientific Research (NWO) Vidi grant 864.14.004 and European Research Council (ERC) Consolidator grant 865694: DiversiPHI. P.C.F. was supported by the Bio-protection Research Centre (Tertiary Education Commission (TEC), New Zealand.

## Author contributions

**Conceptualization**, B.E.D., E.W., P.F. and S.E.M; **Methodology**, A.B., B.E.D., E.W., K.A., P.F., S.E.M and S.M; **Software**, A.B., B.E.D. and K.A.; **Formal analysis**, A.B., B.E.D., K.A. and S.M.; **Investigation**, A.B., B.E.D., K.A., and S.M.; **Data Curation**, A.B., B.E.D., K.A. and S.M.; **Writing – Original draft**, E.W., P.F. and S.M.; **Writing – Review & Editing**, A.B., B.E.D., E.W., K.A., P.F., S.E.M, and S.M.; **Visualization**, S.M.; **Supervision**, B.E.D., E.W., P.F. and S.E.M.; **Project Administration**, B.E.D., E.W., S.E.M. and P.F.; **Funding acquisition**, B.E.D., E.W., S.E.M., S.M. and P.F.

## Supplementary Information

### Supplementary Figures

**Figure S1.**
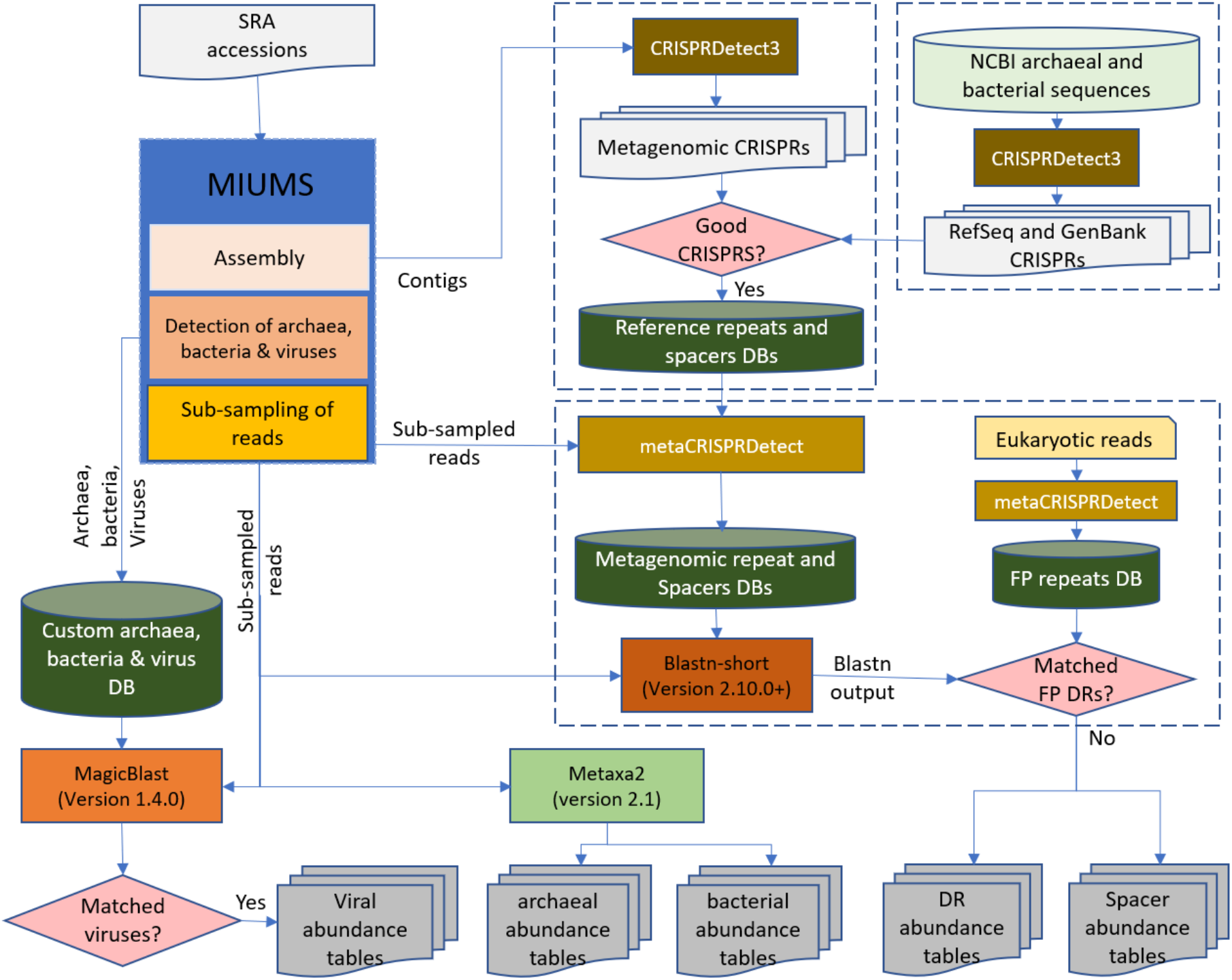
Overview of computational pipeline. Overview of the computational pipeline used to generate CRISPR abundance tables, microbial and viral community abundance tables. In each stage, only the case (yes/no) where data is retained is presented.

**Figure S2.**
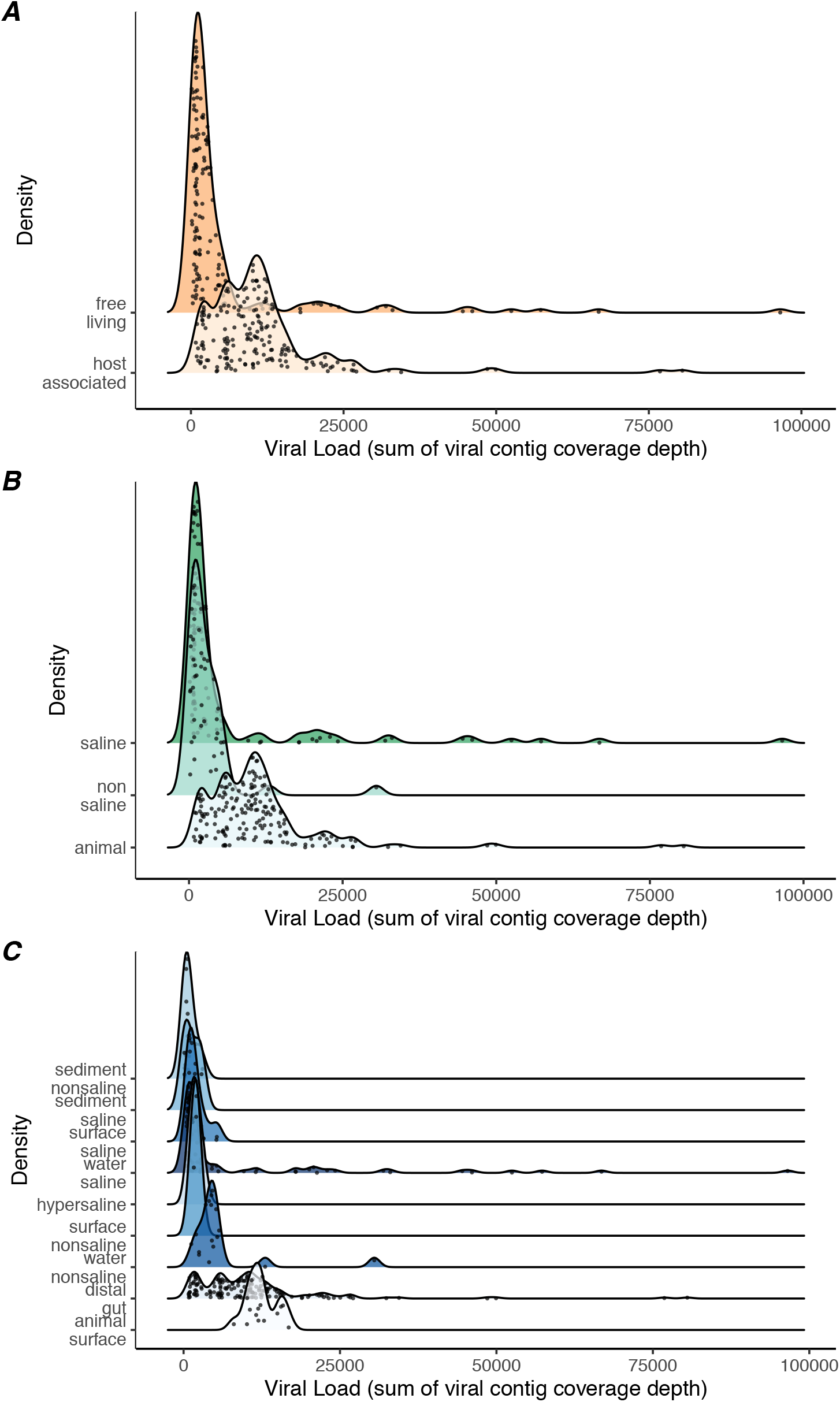
Viral abundance varies by environment. Distributions of viral abundances (sum of viral contig coverage depth per sample) across EMPO level 1 categories (A), level 2 categories (B) and level 3 categories (C). Points represent metagenomic samples.

**Figure S3.**
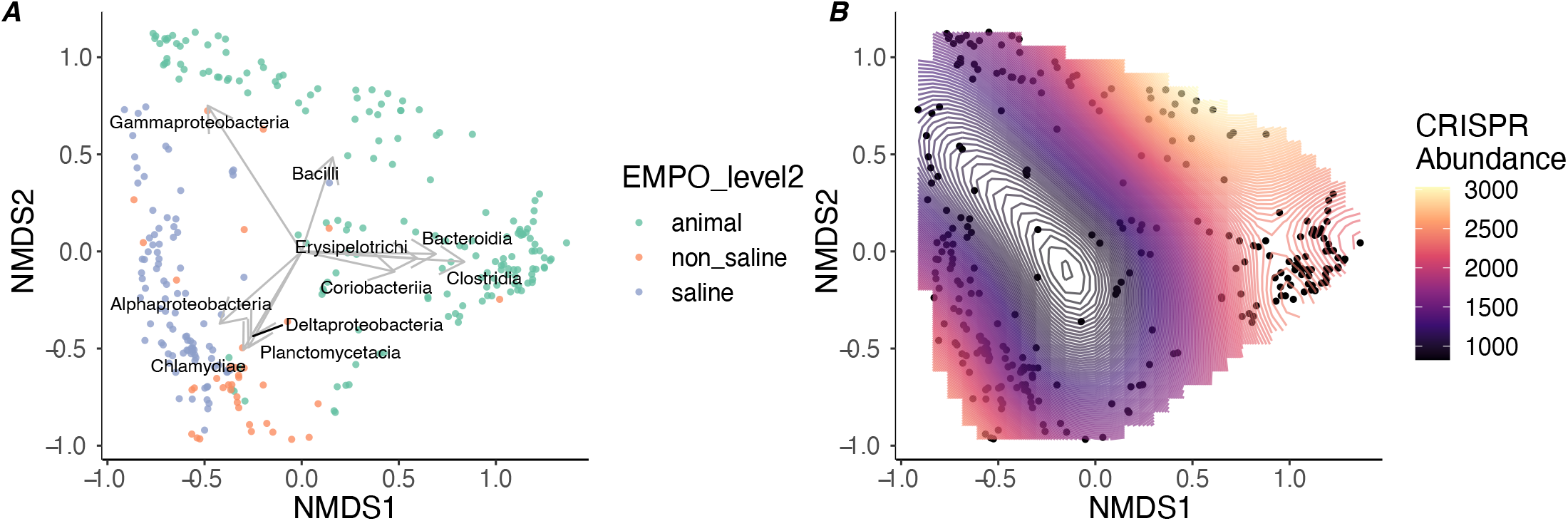
Phylogenetic ordinations of sample community composition. Ordinations of sample community compositions at the Class level. Points represent individual metagenomic samples and are clustered using non-metric multidimensional scaling (NMDS) on Bray-Curtis dissimilarity scores. (*A*) Colors represent the sample classification at EMPO level 2 and arrows represent the top 10 species loadings. (*B*) The same ordination of samples with CRISPR abundance used in a generalized additive model (GAM) to fit a surface predicting CRISPR abundance.

**Figure S4.**
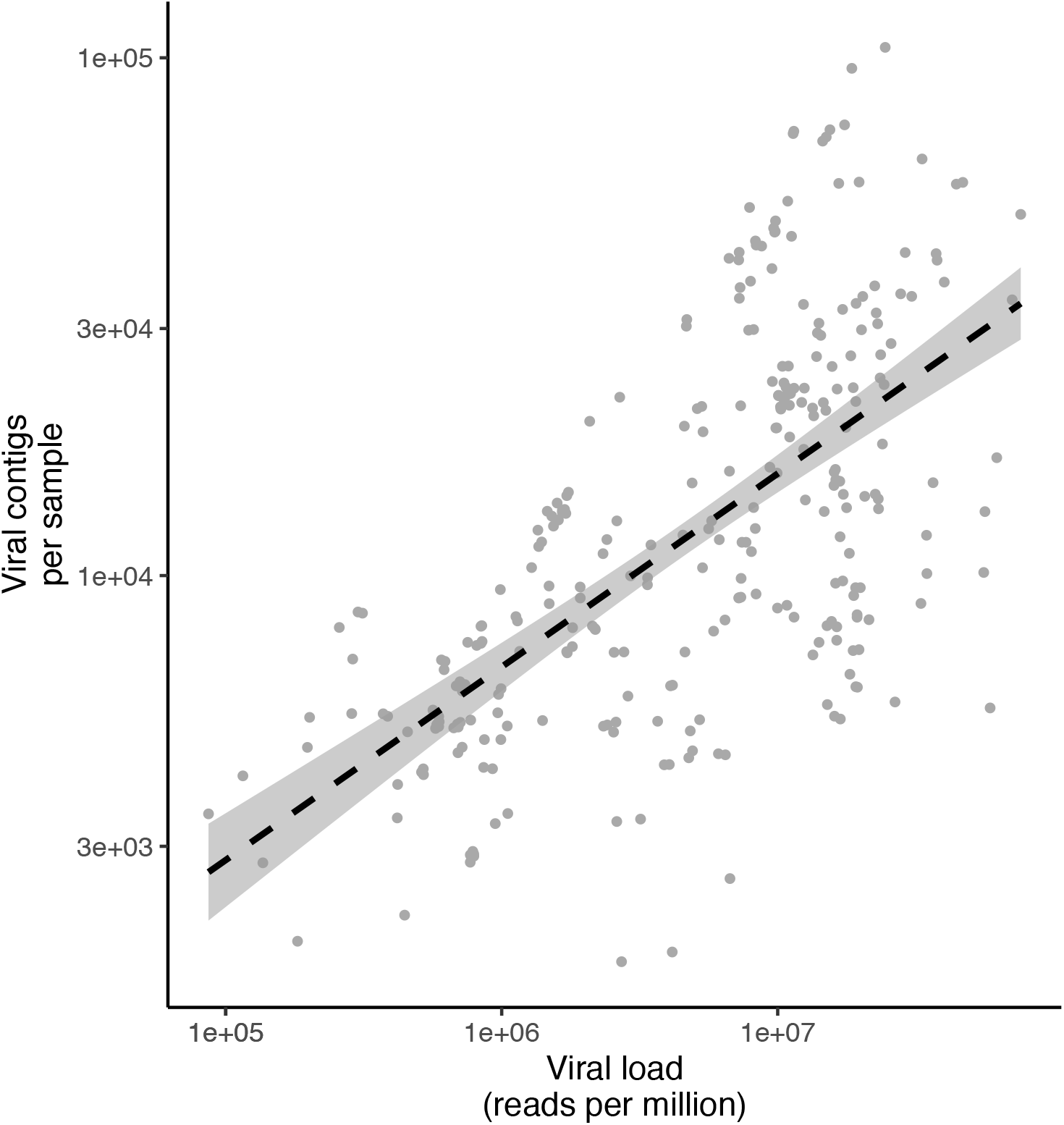
Viral abundance correlates with viral diversity. Correlation between the number of viral contigs per sample and viral abundance (sum of viral contig coverage depth per sample). Line represents linear model fit, shaded areas denote 95% confidence intervals and R^2^ = 0.51.

### Supplementary Tables

**Table S1.** Biome and accession numbers of each metagenome sample. *table_S1_accesions_meta.csv*

**Table S2.** Eukaryotic genome list used for testing false positive rate.

| Species | Genbank accession |
| --- | --- |
| *Triticum aestivum* | GCA_900519105.1 |
| *Drosophila melanogaster* | GCA_000001215.4 |
| *Arabidopsis thaliana* | GCA_000001735.1 |
| *Physcomitrella patens* | GCA_000002425.2 |
| *Caenorhabditis elegans* | GCA_000002985.3 |
| *Heliconius melpomene* | GCA_000313835.1 |

**Table S3.**
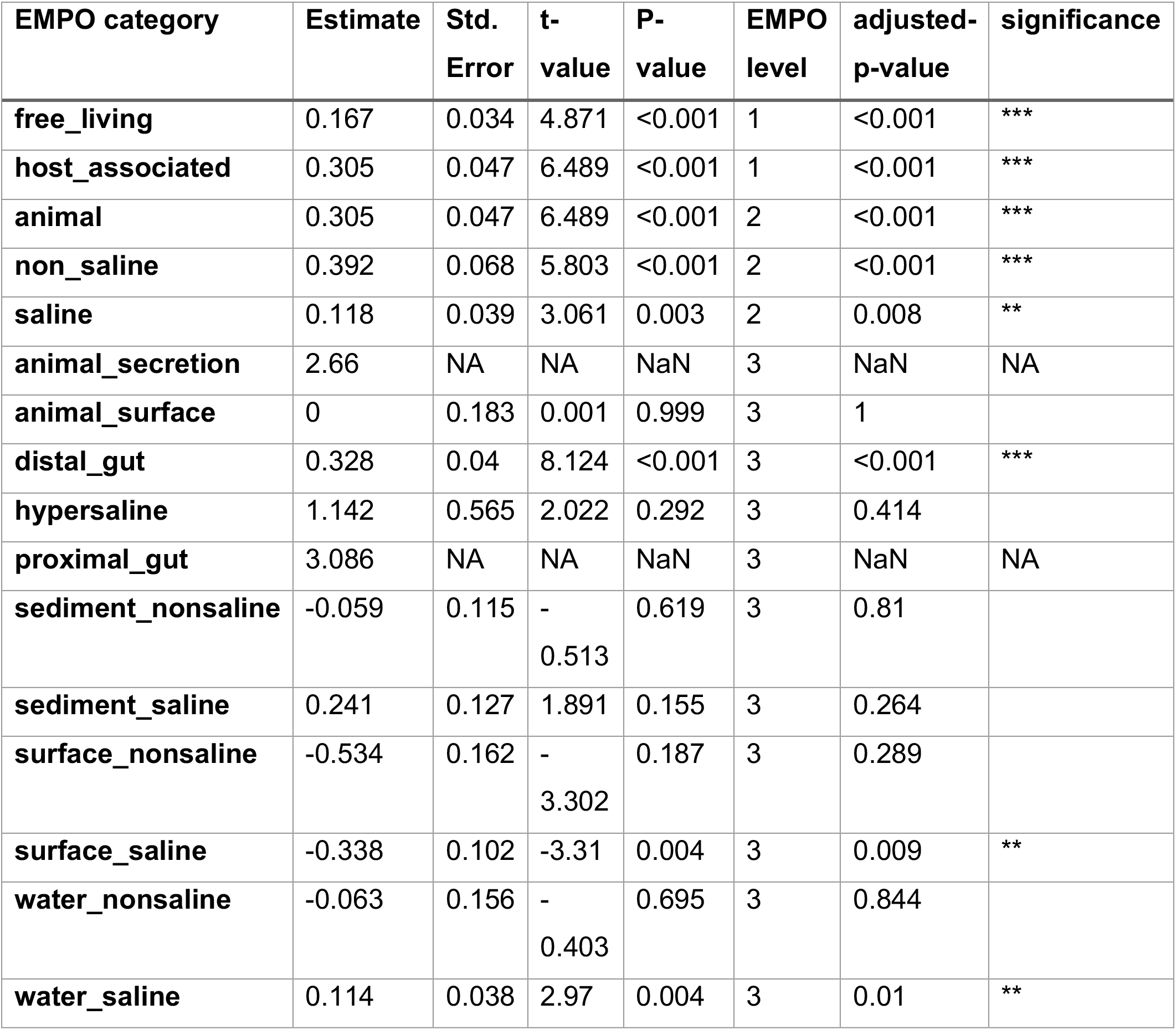
Post-hoc testing of correlations between CRISPR abundance and viral abundance at each environmental classification. Linear regression of CRISPR abundance with viral abundance for each environmental classification. P-values were adjusted using Benjamini-Hochberg correction. NA results are shown for categories with < 3 observations.

| EMPO category | Estimate | Std. Error | t-value | P-value | EMPO level | adjusted-p-value | significance |
| --- | --- | --- | --- | --- | --- | --- | --- |
| free_living | 0.167 | 0.034 | 4.871 | <0.001 | 1 | <0.001 | *** |
| host_associated | 0.305 | 0.047 | 6.489 | <0.001 | 1 | <0.001 | *** |
| animal | 0.305 | 0.047 | 6.489 | <0.001 | 2 | <0.001 | *** |
| non_saline | 0.392 | 0.068 | 5.803 | <0.001 | 2 | <0.001 | *** |
| saline | 0.118 | 0.039 | 3.061 | 0.003 | 2 | 0.008 | ** |
| animal_secretion | 2.66 | NA | NA | NaN | 3 | NaN | NA |
| animal_surface | 0 | 0.183 | 0.001 | 0.999 | 3 | 1 |  |
| distal_gut | 0.328 | 0.04 | 8.124 | <0.001 | 3 | <0.001 | *** |
| hypersaline | 1.142 | 0.565 | 2.022 | 0.292 | 3 | 0.414 |  |
| proximal_gut | 3.086 | NA | NA | NaN | 3 | NaN | NA |
| sediment_nonsaline | -0.059 | 0.115 | -0.513 | 0.619 | 3 | 0.81 |  |
| sediment_saline | 0.241 | 0.127 | 1.891 | 0.155 | 3 | 0.264 |  |
| surface_nonsaline | -0.534 | 0.162 | -3.302 | 0.187 | 3 | 0.289 |  |
| surface_saline | -0.338 | 0.102 | -3.31 | 0.004 | 3 | 0.009 | ** |
| water_nonsaline | -0.063 | 0.156 | -0.403 | 0.695 | 3 | 0.844 |  |
| water_saline | 0.114 | 0.038 | 2.97 | 0.004 | 3 | 0.01 | ** |

**Table S4.** Pearson correlations of CRISPR abundance and viral abundance at each environmental classification.

| **environment** | **EMPO level** | **estimate** | **p.value** | **adjusted.p.value** |
| --- | --- | --- | --- | --- |
| **host associated** | 1 | 0.318669937 | 1.62941E-05 | 0.000179235 |
| **free living** | 1 | 0.225345517 | 0.007648844 | 0.084137289 |
| **animal** | 2 | 0.318669937 | 1.62941E-05 | 0.000179235 |
| **saline** | 2 | 0.279115129 | 0.00588914 | 0.064780536 |
| **non saline** | 2 | 0.300602925 | 0.050152056 | 0.551672619 |
| **distal gut** | 3 | 0.355793262 | 2.37403E-06 | 2.61143E-05 |
| **sediment nonsaline** | 3 | 0.705624736 | 0.003291221 | 0.036203435 |
| **water saline** | 3 | 0.319818959 | 0.006956952 | 0.07652647 |
| **surface saline** | 3 | -0.42738303 | 0.127443734 | 1 |
| **water nonsaline** | 3 | -<br>0.180788442 | 0.487439096 | 1 |
| **animal surface** | 3 | 0.032966836 | 0.907150313 | 1 |

**Table S5.**
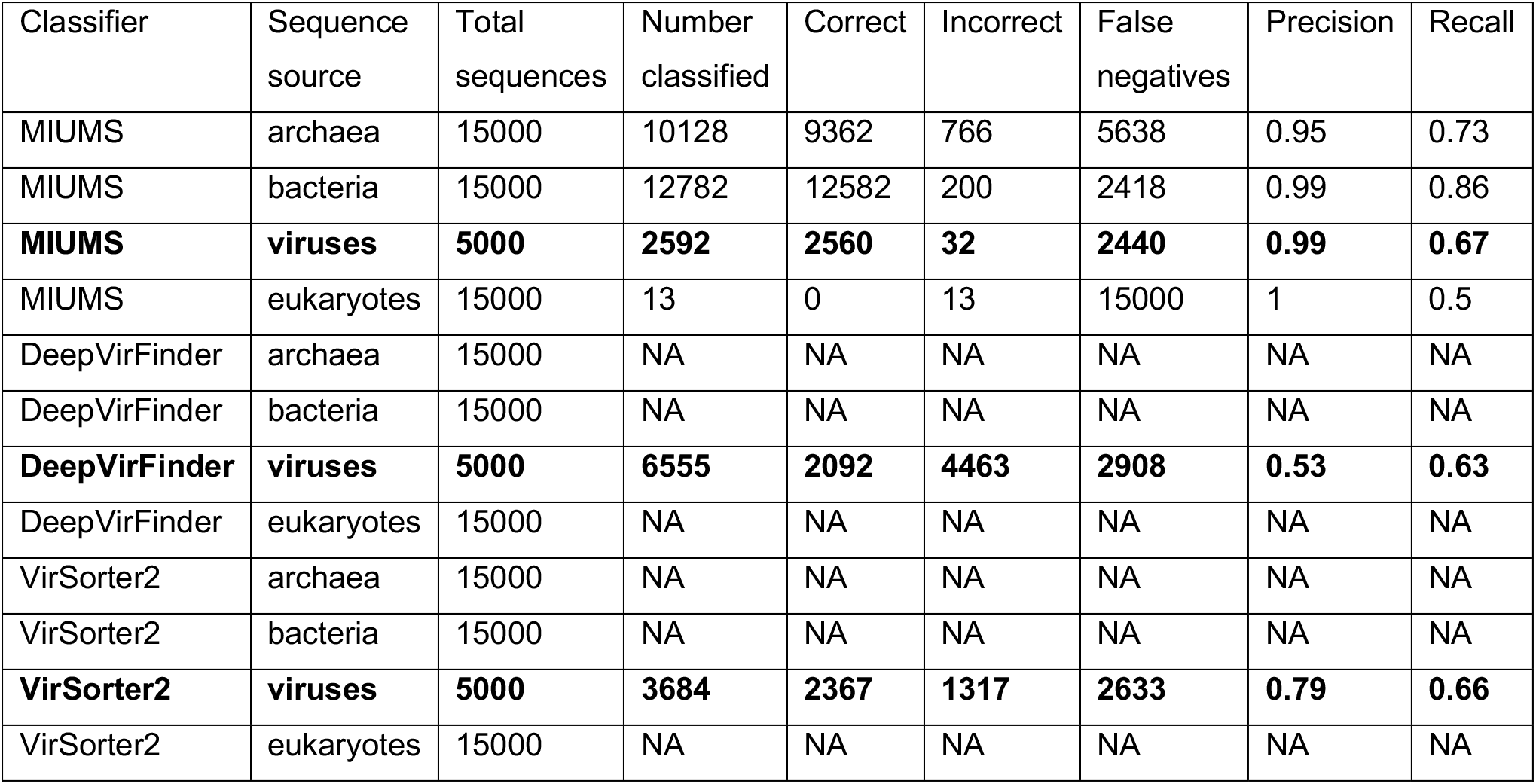
Precision and recall analysis of viral classification tools.

## Supplemental Methods

### Microbial Identification Using Marker Sequence (MIUMS)

MIUMS (version 1.0) utilizes a reference database of protein sequence fragments that are highly specific to their source organism. The process of constructing database is described below.

#### 1. Selection of reference sequences

For the construction of a reference database of marker amino acid sequences, all amino acid sequences from 224 archaeal, 2810 bacterial and 3958 viral (4185) species (published before 1st of March 2017; Refseq release version 79; minimum sequence length of 5000 nucleotides) were selected as the source of the protein sequences for the marker sequence database. These resulted in 155585 archaeal, 2219071 bacterial and 229026 viral amino acid sequences. In addition, a eukaryotic protein sequence database was constructed from 16179736 eukaryotic proteins. The taxonomic information of these protein sequences were collected from NCBI taxonomy files (https://ftp.ncbi.nlm.nih.gov/pub/taxonomy/).

#### 2. Construction of short protein sequence fragments

The construction of a database of marker protein sequence fragments is a multi-step process, which includes i). removal of sequence domains and ii). removal of all potential inter (super)kingdom homologous sequence regions from the target protein sequences.

The selected protein sequences from archaea, bacteria and viruses were screened with Pfam-A HMM profiles (version 30.0) [1] and hmmsearch (HMMER version 3.2.1 with default parameters and --domtblout) [2]. Using the reported domain regions in the sequences, the protein sequences were split into multiple sequence fragments excluding the domain regions.

The sequence fragments were then concatenated into a single protein sequence database and performed an all-vs-all sequence similarity search using diamond (version v0.8.38.100; parameters: --evalue 0.001 --sensitive --no-self-hits) [3]. By analysing the diamond output file, longer sequence fragments which contain shorter sequence fragments were identified and further split into multiple sequence fragments followed by construction of a new sequence database. This process of identification of shorter sequence fragments continued in a cyclic manner till there were no new fragments identified.

#### 3. Removal of inter superkingdom specific protein fragments

The sequence fragments were then separated into their associated superkingdom and screened against the other superkingdom specific primary protein sequences (including eukaryotic protein sequences) using diamond (parameters: --evalue 0.001 --sensitive). Sequence fragments that were found to have inter (super)kingdom matches were identified and removed. This resulted in 610460 archaeal, 8312644 bacterial and 677951 viral marker protein fragments.

#### 4. Assigning taxa specificity to the protein fragments

The archaeal, bacterial, and viral protein fragments were then screened against their own superkingdom specific primary source proteins using diamond (parameters: --evalue 0.001 --sensitive) and the taxa specificity of each protein fragment to each of the higher taxonomic levels (i.e. phylum, class, order, family, genus and species) was determined using lowest common ancestor (LCA) algorithm from all reported diamond matches.

### Taxonomic classification of metagenomic sequences

The default MIUMS runs involve three steps; i) assembly of metagenomic reads, ii) prediction of protein sequence in the assembled contigs using metaGeneMark [4], iii) screening the protein sequences against the marker protein fragment database using diamond (parameters: --evalue 0.001 --sensitive). Contigs with metaGeneMark predicted proteins that contains multiple matches to the protein fragments above stringency cutoffs (e.g. overlapping length >=21aa and sequence identity >=40%) are assigned taxonomy based on the matched protein fragment’s taxa specificity and LCA algorithm. An output table is generated, which shows a list of classified contigs with associated taxonomy.

The raw reads (unless a subsampling of the reads were done) were mapped to the entire assembled contigs using magicblast [5]. The magicblast output file is analysed to identify all reads mapping to the classified contigs with minimum 99% identity and 99% sequence coverage. An output table is generated that shows each of those reads and their associated taxonomy.

### Benchmarking of viral classification tools

In order to assess the accuracy of MIUMS for extracting viral contigs, we compared against existing tools using a test dataset of known sequences. The reference database of marker protein fragments that MIUMS V1.0 uses was constructed from sequences published before 1st of March 2017. Since then the number of newly released viral sequences in the NCBI RefSeq database has nearly doubled (5343 genomic DNA/RNA sequences published between 01-Mar-2017 and 01-June-2021; with minimum length 500 nucleotides). To assess the performance of MIUMS against these newly published sequences; we randomly added 5000 of these viral sequences in a test dataset, comprising closed, whole genomes and contig-level assemblies. The test dataset was also supplemented with 15000 archaea, 15000 bacteria and 15000 eukaryotic sequences published during the same time period mentioned above (with minimum length cut-off of 500 nucleotides and maximum length of 25000 nucleotides). We also included eukaryotic sequences (randomly selected from animal, plant, fungi and protists sequences). This test dataset was analysed with MIUMS V1.0, VirSorter2 (version 2.2.2) [6] and DeepVirFinder (version 1.0) [7] with default parameters. Table S5 shows the summary outputs from the 3 tools used and their respective precision and recall scores.

DeepVirFinder reports a score between 0 to 1 for every input sequence, where a higher score (i.e. close to 1) is a strong indicator of a sequence being viral. Against a minimum score cutoff of 0.95, DeepVirFinder correctly predicted 2092 viruses and falsely predicted 4463 non-viral sequences (Archaea: 1500, Bacteria: 1058, Eukaryotes: 1905) as viruses. While reducing the minimum score-cutoff to 0.75 increases the total number of correctly predicted viral sequences to 3167, it also drastically increases the amount of false predictions to 9944 (Archaea: 3572, Bacteria: 2364, Eukaryotes: 4008). This trend continues with lower minimum score-cutoff to 0.50 (Viruses: 3988, Archaea: 6626, Bacteria: 4417, Eukaryotes: 6678) and 0.25 (Viruses: 4566, Archaea: 9765, Bacteria: 7054, Eukaryotes: 9594).

VirSorter2 generates scores between 0 to 1 (computed on both single and double stranded DNA) and reports potential viral sequences where the maximum of the two scores are >= 0.50. With a minimum score cutoff of 0.95, VirSorter2 correctly predicted 2367 viral sequences with 1317 non-viral sequences (Archaea: 97, Bacteria: 1191, Eukaryotes: 29) falsely predicted as viruses. Reducing the minimum score cutoff to 0.75 increases correctly the predicted viruses to 2798 but increases the falsely predicted non-viral sequences to 2018 (Archaea: 248, Bacteria: 1692, Eukaryotes: 78). At the default score cutoff of 0.50, VirSorter2 correctly predicted 3103 viral sequences with 2427 non-viral sequences (Archaea: 382, Bacteria: 1905, Eukaryotes: 140) falsely predicted to be viral.

In comparison, based on the main taxonomy output table, MIUMS accurately predicted 2557 viruses with a total of 32 non-viral sequences (Archaea: 0, Bacteria: 32, Eukaryotes: 0) falsely predicted to be viruses. The secondary taxonomy output table shows another 540 viral sequences correctly being predicted as potential viruses with 124 non-viral sequences (Archaea: 3, Bacteria: 122, Eukaryote: 0) falsely reported as potential viral sequences. A total of only 13 eukaryotic sequences falsely predicted as archaea, bacteria or viruses (Archaea: 0, bacteria: 12, Viruses: 1).

## References

1. Makarova, K.S., Wolf, Y.I., Iranzo, J., Shmakov, S.A., Alkhnbashi, O.S., Brouns, S.J., Charpentier, E., Cheng, D., Haft, D.H., Horvath, P. and Moineau, S., 2020. Evolutionary classification of CRISPR–Cas systems: a burst of class 2 and derived variants. Nature Reviews Microbiology, 18(2), pp. 67–83

2. Burstein, D., Sun, C.L., Brown, C.T., Sharon, I., Anantharaman, K., Probst, A.J., Thomas, B.C. and Banfield, J.F., 2016. Major bacterial lineages are essentially devoid of CRISPR-Cas viral defence systems. Nature communications, 7(1), pp. 1–8.

3. Pourcel, C., Touchon, M., Villeriot, N., Vernadet, J.P., Couvin, D., Toffano-Nioche, C. and Vergnaud, G., 2020. CRISPRCasdb a successor of CRISPRdb containing CRISPR arrays and cas genes from complete genome sequences, and tools to download and query lists of repeats and spacers. Nucleic acids research, 48(D1), pp.D535-D544.

4. Mojica, F.J. and Garrett, R.A., 2013. Discovery and seminal developments in the CRISPR field. In CRISPR-Cas Systems (pp. 1–31). Springer, Berlin, Heidelber

5. Makarova, K.S., Wolf, Y.I. and Koonin, E.V., 2013. Comparative genomics of defense systems in archaea and bacteria. Nucleic acids research, 41(8), pp. 4360–4377.

6. Hille, F., Richter, H., Wong, S.P., Bratovič, M., Ressel, S. and Charpentier, E., 2018. The biology of CRISPR-Cas: backward and forward. Cell, 172(6), pp. 1239–1259.

7. Gurney, J., Pleška, M. and Levin, B.R., 2019. Why put up with immunity when there is resistance: an excursion into the population and evolutionary dynamics of restriction–modification and CRISPR-Cas. Philosophical Transactions of the Royal Society B, 374(1772), p. 20180096.

8. Iranzo, J., Lobkovsky, A.E., Wolf, Y.I. and Koonin, E.V., 2013. Evolutionary dynamics of the prokaryotic adaptive immunity system CRISPR-Cas in an explicit ecological context. Journal of bacteriology, 195(17), pp. 3834–3844.

9. Weinberger, A.D., Sun, C.L., Pluciński, M.M., Denef, V.J., Thomas, B.C., Horvath, P., Barrangou, R., Gilmore, M.S., Getz, W.M. and Banfield, J.F., 2012. Persisting viral sequences shape microbial CRISPR-based immunity. PLoS Comput Biol, 8(4), p. e1002475.

10. Westra, E.R., van Houte, S., Oyesiku-Blakemore, S., Makin, B., Broniewski, J.M., Best, A., Bondy-Denomy, J., Davidson, A., Boots, M. and Buckling, A., 2015. Parasite exposure drives selective evolution of constitutive versus inducible defense. Current biology, 25(8), pp. 1043–1049.

11. Broniewski, J.M., Meaden, S., Paterson, S., Buckling, A. and Westra, E.R., 2020. The effect of phage genetic diversity on bacterial resistance evolution. The ISME journal, 14(3), pp. 828–836.

12. Westra, E.R. and Levin, B.R., 2020. It is unclear how important CRISPR-Cas systems are for protecting natural populations of bacteria against infections by mobile genetic elements. Proceedings of the National Academy of Sciences, 117(45), pp. 27777–27785.

13. Weissman, J.L., Laljani, R.M., Fagan, W.F. and Johnson, P.L., 2019. Visualization and prediction of CRISPR incidence in microbial trait-space to identify drivers of antiviral immune strategy. The ISME journal, 13(10), pp. 2589–2602.

14. Thompson, L.R., Sanders, J.G., McDonald, D., Amir, A., Ladau, J., Locey, K.J., Prill, R.J., Tripathi, A., Gibbons, S.M., Ackermann, G. and Navas-Molina, J.A., 2017. A communal catalogue reveals Earth’s multiscale microbial diversity. Nature, 551(7681), pp. 457–463.

15. Wu, R., Chai, B., Cole, J.R., Gunturu, S.K., Guo, X., Tian, R., Gu, J.D., Zhou, J. and Tiedje, J.M., 2020. Targeted assemblies of cas1 suggest CRISPR-Cas’s response to soil warming. The ISME journal, pp.1-12.

16. Münch, P.C., Franzosa, E.A., Stecher, B., McHardy, A.C. and Huttenhower, C., 2021. Identification of Natural CRISPR Systems and Targets in the Human Microbiome. Cell Host & Microbe, 29(1), pp. 94–106.

17. Bernheim, A. and Sorek, R., 2020. The pan-immune system of bacteria: antiviral defence as a community resource. Nature Reviews Microbiology, 18(2), pp. 113–119.

18. Bernheim, A., Calvo-Villamañán, A., Basier, C., Cui, L., Rocha, E.P., Touchon, M. and Bikard, D., 2017. Inhibition of NHEJ repair by type II-A CRISPR-Cas systems in bacteria. Nature communications, 8(1), pp. 1–9.

19. Hernandez, C.A. and Koskella, B., 2019. Phage resistance evolution in vitro is not reflective of in vivo outcome in a plant-bacteria-phage system. Evolution, 73(12), pp. 2461–2475.

20. Laanto, E., Bamford, J.K., Laakso, J. and Sundberg, L.R., 2012. Phage-driven loss of virulence in a fish pathogenic bacterium. PLoS One, 7(12), p. e53157.

21. Alseth, E.O., Pursey, E., Luján, A.M., McLeod, I., Rollie, C. and Westra, E.R., 2019. Bacterial biodiversity drives the evolution of CRISPR-based phage resistance. Nature, 574(7779), pp. 549–552.

22. Smith, L.M., Jackson, S.A., Malone, L.M., Ussher, J.E., Gardner, P.P. and Fineran, P.C., 2021. The Rcs stress response inversely controls surface and CRISPR–Cas adaptive immunity to discriminate plasmids and phages. Nature crobiology, 6(2), pp. 162–172.

23. Arbas, S.M., Narayanasamy, S., Herold, M., Lebrun, L.A., Hoopmann, M.R., Li, S., Lam, T.J., Kunath, B.J., Hicks, N.D., Liu, C.M. and Price, L.B., 2021. Roles of bacteriophages, plasmids and CRISPR immunity in microbial community dynamics revealed using time-series integrated meta-omics. Nature microbiology, 6(1), pp. 123–135.

24. Watson, B.N., Steens, J.A., Staals, R.H., Westra, E.R. and van Houte, S., 2021. Coevolution between bacterial CRISPR-Cas systems and their bacteriophages. Cell Host & Microbe, 29(5), pp. 715–725.

25. Puigbò, P., Makarova, K.S., Kristensen, D.M., Wolf, Y.I. and Koonin, E.V., 2017. Reconstruction of the evolution of microbial defense systems. BMC evolutionary biology, 17(1), pp. 1–13.

26. van Belkum, A., Soriaga, L.B., LaFave, M.C., Akella, S., Veyrieras, J.B., Barbu, E.M., Shortridge, D., Blanc, B., Hannum, G., Zambardi, G. and Miller, K., 2015. Phylogenetic distribution of CRISPR-Cas systems in antibiotic-resistant Pseudomonas aeruginosa. MBio, 6(6), pp. e01796–15.

27. Watson, B.N., Staals, R.H. and Fineran, P.C., 2018. CRISPR-Cas-mediated phage resistance enhances horizontal gene transfer by transduction. MBio, 9(1), pp. e02406–17.

28. Varble, A., Meaden, S., Barrangou, R., Westra, E.R. and Marraffini, L.A., 2019. Recombination between phages and CRISPR-cas loci facilitates horizontal gene transfer in staphylococci. Nature microbiology, 4(6), pp. 956–963.

29. Bushnell, B., Rood, J. and Singer, E., 2017. BBMerge–Accurate paired shotgun read merging via overlap. PloS one, 12(10), p. e0185056.

30. Li, D., Liu, C.M., Luo, R., Sadakane, K. and Lam, T.W., 2015. MEGAHIT: an ultra-fast single-node solution for large and complex metagenomics assembly via succinct de Bruijn graph. Bioinformatics, 31(10), pp. 1674–1676.

31. Bengtsson-Palme, J., Hartmann, M., Eriksson, K.M., Pal, C., Thorell, K., Larsson, D.G.J. and Nilsson, R.H., 2015. METAXA2: improved identification and taxonomic classification of small and large subunit rRNA in metagenomic data. Molecular ecology resources, 15(6), pp. 1403–1414.

32. Boratyn, G.M., Thierry-Mieg, J., Thierry-Mieg, D., Busby, B. and Madden, T.L., 2019. Magic-BLAST, an accurate RNA-seq aligner for long and short reads. BMC bioinformatics, 20(1), pp. 1–19.

33. Biswas, A., Staals, R.H., Morales, S.E., Fineran, P.C. and Brown, C.M., 2016. CRISPRDetect: a flexible algorithm to define CRISPR arrays. BMC genomics, 17(1), pp. 1–14.

34. Li, W. and Godzik, A., 2006. Cd-hit: a fast program for clustering and comparing large sets of protein or nucleotide sequences. Bioinformatics, 22(13), pp. 1658–1659.

35. Chen, Y., Ye, W., Zhang, Y. and Xu, Y., 2015. High speed BLASTN: an accelerated MegaBLAST search tool. Nucleic acids research, 43(16), pp. 7762–7768.

36. Altschul, S.F., Gish, W., Miller, W., Myers, E.W. and Lipman, D.J., 1990. Basic local alignment search tool. Journal of molecular biology, 215(3), pp. 403–410.

37. Li, H. and Durbin, R., 2009. Fast and accurate short read alignment with Burrows–Wheeler transform. bioinformatics, 25(14), pp. 1754–1760.

38. Nei, M., 1978. Estimation of average heterozygosity and genetic distance from a small number of individuals. Genetics, 89(3), pp. 583–590.

39. Koboldt, D.C., Chen, K., Wylie, T., Larson, D.E., McLellan, M.D., Mardis, E.R., Weinstock, G.M., Wilson, R.K. and Ding, L., 2009. VarScan: variant detection in massively parallel sequencing of individual and pooled samples. Bioinformatics, 25(17), pp. 2283–2285.

40. Kang, D.D., Froula, J., Egan, R. and Wang, Z., 2015. MetaBAT, an efficient tool for accurately reconstructing single genomes from complex microbial communities. PeerJ, 3, p. e1165.

41. Oksanen, J., Blanchet, F.G., Kindt, R., Legendre, P., Minchin, P.R., O’hara, R.B., Simpson, G.L., Solymos, P., Stevens, M.H.H. and Wagner, H., 2013. Community ecology package. R package version, 2(0)

## References

1. Finn RD, Bateman A, Clements J, Coggill P, Eberhardt RY, Eddy SR, et al. Pfam: the protein families database. Nucleic Acids Res 2014; 42: D222–30.

2. Eddy SR. Profile hidden Markov models. Bioinformatics. 1998. Oxford University Press., 14: 755–763

3. Buchfink B, Xie C, Huson DH. Fast and sensitive protein alignment using DIAMOND. Nat Methods. 2014. Nature Publishing Group., 12: 59–60

4. Zhu W, Lomsadze A, Borodovsky M. Ab initio gene identification in metagenomic sequences. Nucleic Acids Res 2010; 38: e132–e132.

5. Boratyn GM, Thierry-Mieg J, Thierry-Mieg D, Busby B, Madden TL. Magic-BLAST, an accurate RNA-seq aligner for long and short reads. BMC Bioinformatics 2019; 20: 405.

6. Guo J, Bolduc B, Zayed AA, Varsani A, Dominguez-Huerta G, Delmont TO, Pratama AA, Gazitúa MC, Vik D, Sullivan MB, Roux S. VirSorter2: a multi-classifier, expert-guided approach to detect diverse DNA and RNA viruses. Microbiome. 2021 Dec; 9:1–3.

7. Ren J, Song K, Deng C, Ahlgren NA, Fuhrman JA, Li Y, Xie X, Poplin R, Sun F. Identifying viruses from metagenomic data using deep learning. Quantitative Biology. 2020 Jan 22:1–4.

